# Crosstalk between Immune Microenvironment and Hair Follicle Cells Underlies Sexual Dimorphism in Androgenetic Alopecia

**DOI:** 10.1101/2025.09.28.678997

**Authors:** Feifei Du, Lei Ge, Yali Yang, Jiaming Wang, Zichen Sun, Jie Chen, Xiameng Li, Tianxia Xiao, Zhili Deng, Dafu Zhi, Jian V Zhang

**Affiliations:** Center for Energy Metabolism and Reproduction, Shenzhen Institutes of Advanced Technology, Chinese Academy of Sciences, Shenzhen, Guangdong, 518055, China; Shenzhen Key Laboratory of Metabolic Health, Shenzhen Metabolism and Reproductive Targeted Delivery Proof-of-Concept Center, Shenzhen Institutes of Advanced Technology, Chinese Academy of Sciences, Shenzhen, Guangdong, 518055, China; Health Sciences Institute, China Medical University, Shenyang 110122, China; Department of Biomedical Sciences, Faculty of Health Sciences, University of Macau, Macau, 999078, China; Department of Dermatology, Xiangya Hospital, Central South University, Changsha, 410008, Hunan, China; Hunan Key Laboratory of Aging Biology, Xiangya Hospital, Central South University, Changsha, 410008, Hunan, China; Zhongke Haohanhang Biotechnology (Shen zhen) Co., Ltd, Shenzhen, Guangdong, 51800, China; Faculty of Pharmaceutical Sciences, Shenzhen University of Advanced Technology, Shenzhen, Guangdong, 518028, China; Sino-European Center of Biomedicine and Health, Shenzhen, Guangdong, 518000, China

**Keywords:** Androgenetic alopecia, dihydrotestosterone, dermal papilla cells, skin-resident myeloid cells, Y001, Y002

## Abstract

Androgenetic alopecia (AGA), also known as female pattern hair loss (FPHL) in women, is the most prevalent form of hair loss. It is characterized by progressive miniaturization of hair follicles and shortening of the anagen phase. The condition is widely attributed to genetic predisposition and androgen-mediated activation of androgen receptors. Other factors, such as the immune microenvironment, could also contribute to the pathogenesis. However, the specific mechanisms involved are still poorly understood. This study aimed to investigate the potential role of skin-resident myeloid cells in interacting with hair follicle cells under androgen stimulation, and to elucidate the sex-specific differences in dihydrotestosterone (DHT)-induced hair loss. Both female and male mice received DHT treatment, and histological examination was performed to assess DHT-induced alterations in skin morphology. Single-cell RNA sequencing (scRNA-seq) was utilized to profile skin cell populations and explore underlying mechanisms. Our results demonstrated that DHT inhibited hair regrowth in both sexes, altered skin thickness, and induced hair follicle miniaturization. scRNA-seq analysis revealed enhanced interactions between myeloid and fibroblast subpopulations, with more robust crosstalk observed in female mice. An in vitro experiment demonstrated that DHT promoted apoptosis of dermal papilla cells (DPCs) in the presence of macrophages. Treatment with polypeptides Y001 and Y002 effectively promoted hair regrowth by suppressing apoptosis signaling pathways. Our findings underscore the interactions between immune cells and hair follicular cells, particularly mediated by skin-resident myeloid cells, in the sexual dimorphism of androgenetic alopecia. The polypeptides Y001 and Y002 exhibit promising therapeutic potential by targeting apoptotic pathways, offering novel avenues for AGA treatment.

## Introduction

Androgenetic alopecia (AGA) is a common form of hair loss that affects both men and women. In women, this condition is specifically referred to as female pattern hair loss (FPHL). It is estimated that more than 50% men at the age of 50 are affected by this condition, and its prevalence continues to increase with age[1, 2]. In contrast, the incidence in women of the same age is gengerally lower, however, the incidence in women follows a bimodal distribution, with peaks occurring during reproductive years and menopause, and continues to rise with age, reaching approximately 55% of women over 70.[3, 4]. Ethnicity is another risk factor. A study conducted across Chinese six-cities reported that the prevalence of AGA among Asian populations is significantly lower than in Caucasians, for both men and women[5]. Surprisingly, there has been an increasing trend of AGA occurring among adolescents and even pediatric populations[6]. To date, only two treatments have been approved officially by the US Food and Drug Administration (FDA) for this type of hair loss. However, due to drawbacks such as significant side effects and high relapse rates, neither is considered fully satisfactory[7]. Other therapies have also demonstrated some efficacy but remain insufficiently effective[7, 8].

Androgenetic alopecia (AGA) is a progressive hair loss disorder involving follicular miniaturization and a shortened anagen phase. Sexual dimorphism has been observed in various AGA patients, which is reflected not only in incidence rates but also in treatment efficacy[9, 10], hormonal profiles[11], and genetic susceptibility[12]. Clinically, AGA manifests differently between sexes. In male, AGA is classified by using the Hamilton–Norwood scale, typically presenting with bitemporal recession and vertex thinning, whereas female pattern hair loss follows the Ludwig system, more commonly exhibiting diffuse crown thinning with preserved frontal hairline[13]. Serum androgen profiles differ as well. Men with AGA frequently demonstrate elevated systemic androgen levels[14, 15]. In contrast, findings in women are inconsistent, some studies reported increased levels of specific metabolites, such as 3α-androstanediol and 3α-androstanediol sulfate[16], while others show no significant difference compared to unaffected women[17]. This divergence may stem from the sexual differences of key enzymes expression in hair follicles, including 5α-reductase types I and II, androgen receptor, and cytochrome P450 aromatase, resulting in distinct local hormonal microenvironments between sexes[18]. Genetic studies further highlight sex-based differences. Genome-wide association studies have identified sex-specific susceptibility loci and single-nucleotide polymorphisms (SNPs)[12, 19]. Additional factors, such as microinflammation, angiogenesis, and follicular stem cell regeneration have also been implicated in AGA[20-22]. However, whether their contributions vary by sex, and hence help explain the differential pathogenesis and progression between males and females, remain an open question worthy of further investigation.

Emerging research highlights a correlation between the immune microenvironment and the progression of AGA[23]. In a study of 58 FPHL specimens, perifollicular lymphocytic infiltration in the majority of samples, particularly in the infundibular and isthmic regions was observed.

Notably, inflammatory cells were present adjacent to miniaturized follicles in 86.2% of cases[24]. Supporting this finding, another investigation reported perifollicular fibrosis in the lower infundibulum and isthmus in approximately 76% of bald scalp samples from AGA patients. Moreover, moderate to dense perifollicular lymphocytic infiltrates were detected in about 33% of patients, whereas no such inflammation was observed in control subjects[25]. A cross-sectional study, involving 17 FPHL patients, further revealed modest mononuclear infiltration around hair follicles, with a positive correlation between inflammation and apoptosis[26]. In addition, it has been proven that overweight, a condition linked to chronic inflammation, has been strongly associated with early-onset AGA in men[27]. Obesity-mediated stress could accelerate hair follicle miniaturization and finally hair thinning, potentially through mechanisms involving inflammaging of stem cells and pro-inflammatory signaling within follicular niches[28]. Growing evidence have indicated that anti-inflammatory treatments may ameliorate AGA conditions, although further studies are needed to confirm their efficacy and safety [29, 30].

In this study, we investigated the mechanisms of androgen-induced alopecia by using both male and female mouse models, with a specific emphasis on identifying sex-related differences in pathogenesis. A particular focus was placed on cell-cell interactions between skin-resident myeloid cells and fibroblast cells. Furthermore, we evaluated two polypeptides as novel therapeutic interventions for AGA in mouse models, both of which demonstrated promising efficacy in promoting hair regrowth in mice of both sexes.

## Methods

### Mice

All animal experiments were approved by the Institutional Animal Care and Use Committee (IACUC, Permit number: APS-230217-003-01). C57BL/6J were purchased from the Charles River Laboratories (Guangdong, China) and acclimatized for 1 week prior to experimentation. All mice were housed under specific pathogen-free conditions with a 12-hour light/dark cycle and provided ad libitum access to food and water.

### DHT-induced alopecia mouse model

The mice of each gender were randomly assigned to two groups, with 3-4 mice for each group. 7 weeks old mice were intraperitoneal injected either 2mg/kg/d DHT (dissolved in corn oil, Solarbio, Beijing, China) or vehicle (control group) every other day. The dorsal skin of all mice was depilated using shaving followed by hair removal cream. All mice were anesthetized with 2.5% Avertin and kept warm during the procedure of depilation and photographing. Hair regrowth was subsequently monitored and recorded every other day for a period of 24 days. To study the role of Y001 (PCT/CN2024/080241; CN2024102020323) and Y002 (PCT/CN2024/079940; CN2024102098116) on hair regrowth process, both female and male mice were topically administrated Y001 (0.02 mg/g/d for female mice, 0.08mg/g/d for male mice) or Y002 (0.08 mg/g/d) daily. Mice were anesthetized followed by cervical neck dislocation prior to dorsal skin collection.

### Hematoxylin and eosin (H&E) staining

The dorsal skin samples were collected from female mice at day 10 or male mice at day 12 after depilation. Tissues were then fixed in 4% paraformaldehyde, embeded in paraffin, and sectioned in 5μm. The sections were deparaffinized and rehydrated through a graded ethanol series to distilled water, and then stained with hematoxylin, followed by washing under running tap water. The sections were differentiated in 1% acid alcohol (1% HCl in ethanol) for 1–3 seconds and rinsed again under running tap water. Cytoplasmic counterstaining was performed using eosin. Dehydration was carried out through three changes of 95% ethanol and two changes of 100% ethanol. Tissues were cleared in three changes of xylene, and finally mounted with a coverslip using resinous mounting medium. Images were acquired using a light microscope (Olympus, Tokyo, Japan). The diameters of hair follicles and dermal thickness ratio were quantified using ImageJ software.

### Masson trichrome staining

Masson trichrome staining was conducted according to the manufactures’ instruction (G1340, Solarbio, Beijing, China). Briefly, skin sections were deparaffinized and hydrated, then stained with weigert’s iron hematoxylin solution for 5min. Differentiated in acid-alcohol solution for 5s, the slides were blued in bluing solution for 3min. Aeftr rinsing for 1min, the slides were then stained with ponceau-acid fucshin solution for 5min and differentiated in phosphomolybic acid solution for 1min. Subsequently, sections were incubated with aniline bule solution for 1min. treated with acetic acid working solution for 1 minute, and dehydrated through a graded ethanol series (95% and absolute ethanol, three times each). Finally, the slides were mounted with resinene. Images were acquired using a light microscope (Olympus, Tokyo, Japan) and analyzed with ImageJ software.

### Immunohistochemistry (IHC)

Skin sections were deparaffinized and rehydrated firstly, then the sections were boiled in citrate buffer by microwave oven. Quench slides with 3% hydrogen peroxide (H_2_O_2_) solution for 30min, and block the slides with 1% BSA for 1h. Incubated with anti-androgen receptor antibody (abcam, Waltham, MA, USA) overnight at −4°C. Second antibody was added the next day and incubated for 30min, add diaminobenzidine (DAB) substrate to each slide and wait for appropriate color development and then stopped with dH_2_O. After that, the slides were counterstained with hematoxylin for 3min, differentiated in acid-alcohol solution, and blued in bluing solution for 3min. Then the slides were hydrated and sealed with resinene. The images were taken by microscope (Olympus, Tokyo, Japan) and analyzed by ImageJ.

### Single cell suspension preparation and sequencing

Skin samples were chopped into small pieces and digested at 37°C for about 30min. After digestion, the cell suspension was centrifuged, and the supernatant was discarded. Cells were resuspended with 1 mL of 1640 containing 2% FBS. Then cells were filtered with a 30μm-cell-strainer. Lastly, the concentration of the cell suspension was adjusted to 700-1200 cells/uL. Then cells were loaded into microfluidic chip of Chip A Single Cell Kit v2.1 (MobiDrop (Zhejiang) Ltd., cat. no. S050100301) to generate droplets with MobiNova-100(MobiDrop (Zhejiang) Ltd., cat. no.A1A40001). Each cell was involved into a droplet which contained a gel bead linked with up to millions oligos (cell unique barcode). After encapsulation, droplets suffer light cut by MobiNovaSP-100(MobiDrop (Zhejiang) Ltd., cat. no.A2A40001) while oligos diffuse into reaction mix. The mRNAs were captured by cell barcodes with cDNA amplification in droplets. Following reverse transcription, cDNAs with barcodes were amplified, and a library was constructed using the High Throughput Single-Cell 3’ Transcriptome Kit v2.1 (MobiDrop (Zhejiang) Co., Ltd., cat. no. S050200301) and the 3’ Dual Index Kit(MobiDrop (Zhejiang) Ltd., cat. no. S050300301). The resulting libraries were sequenced on an Illumina System according to the manufacturer’s instructions(Illumina) at Chi Biotech Ltd.

### Single cell data analysis

Fastp was used to perform basic statistics on the quality of the raw reads. Raw datas (fastq format) of single-cell 3’ transcriptome were pre-analyzed by MobiVision (version 3.2, MobiDrop), Filtered cell-gene matrix was obtained with MobiVision. The Seurat package was used to normalize data, dimensionality reduction, clustering, differential expression. Seurat alignment method canonical correlation analysis (CCA) was used for integrated analysis of datasets. For clustering, highly variable genes were selected and the principal components based on those genes used to build a graph. Gene Ontology (GO) and KEGG enrichment analysis of marker genes was implemented by the clusterProfiler R package, in which gene length bias was corrected. Terms with corrected Pvalue less than 0.05 were considered significantly enriched by marker genes.

### Bulk RNA sequencing and analysis

Topical interventions with Y001 and Y002 were administered on DHT-induced alopecia mouse models. The dorsal skin samples were haversted at indicated time points for bulk RNA sequencing. Total RNA was extracted by using TRIzol (Invitrogen, California, USA), and the samples were subsequently submitted to OE Biotechnology Ltd for sequencing (Shanghai, China). Library construction and sequencing were performed on an Illumina NovaSeq 6000 platform following the company’s standard operational procedures. Gene expression levels were quantified using the FPKM for visualization. Differentially expressed genes (DEGs) were carried out with DESeq2 (version 1.22.2) based on raw counts, with Benjamini–Hochberg (BH) FDR correction applied to adjust p-values. Genes with an p-value < 0.05 and |log2 fold change| > 1 were considered significantly differentially expressed. Gene Set Enrichment Analysis (GSEA) was performed based on a predefined gene set of FPKM values. Multiple testing correction was applied using the default FDR method within the GSEA algorithm.

### Cell Culture and Coculture Experiments

Immortalized human dermal papilla cells (hDPCs) were generated as previously decribed[21], which was provided by professor Deng, and maintained in Dulbecco’s Modified Eagle Medium (DMEM) supplemented with 10% fetal bovine serum, 1% penicillin and streptomycin. For coculture experiments, serum-starved hDPCs were seeded into the lower chamber of a 24-well Transwell system (0.4 μm pore size; Corning Inc., Corning, NY) at a density of 4 × 10^5^ cells per well. RAW264.7 macrophages (2 × 10^4^ cells per well) were plated in the upper chamber. The following experimental groups were established: hDPCs alone or hDPCs cocultured with macrophages, each treated with or without 100 μM dihydrotestosterone (DHT, Solarbio, China) for 24 hours. After treatment, hDPCs were collected for subsequent analysis.

### Flow cytometry

To assess DHT-induced apoptosis, hDPCs were analyzed using an Annexin V-FITC/PI apoptosis detection kit (A211, Vazyme, China) according to the manufacturers’ instructions. Briefly, hDPCs from cocultured experiment were harvested by using EDTA-free trypsin, washed twice with cold PBS, and centrifuged. The pellet was resuspended in 100 µl of 1× binding buffer. Then, 5 µl of Annexin V-FITC and 5 µl of propidium iodide (PI) staining solution were added to each sample and incubated for 10 minutes in the dark. Subsequently, 400 µl of 1× binding buffer was added to each tube. Samples were loaded immediately to a CytoFLEX flow cytometer (Beckman Coulter, Indianapolis, IN, USA) and data was processed with FlowJo 10.9. The total apoptosis rate was defined as the sum of early (Annexin V^+^/PI^−^) and late (Annexin V^+^/PI^+^) apoptotic cells.

### Quantitative real-time Polymerase Chain Reaction

Total RNA was isolated by Trizol reagent (Invitrogen, California, USA), the concentration was determined by by measuring the absorbance at 260 nm using a NanoDrop spectrophotometer (Thermo Fisher Scientific, MA, USA). cDNA synthesis was conducted by PrimeScript RT Master Mix (Takara Bio, Shiga, Japan) in a volume of 20 μl according to the manufacturer’s instructions. Quantitative real-time PCR was performed with SYBR qPCR Mix (Toyobo, Osaka, Japan) in a final volume of 20 μl. The primer sequences of β-actin, IGF1R, and Bax were as follows:

β-actin: Forward: 5’-AGCGAGCATCCCCCAAAGTT-3’,

Reverse:5’-GGGCACGAAGGCTCATCATT-3’;

IGF1R: Forward:5’-GGACAGGTCAGAGGGTTTC-3’,

Reverse: 5’-CTCGTAACTCTTCTCTGTGCC-3’;

Bax: Forward: 5’-TTTTGCTTCAGGGTTTCATCCA-3’,

Reverse: 5’-TGCCACTCGGAAAAAGACCTC-3’;

SDC1: Forward: 5’-ACGGCTATTCCCACGTCTC-3’

Reverse : 5’-TCTGGCAGGACTACAGCCTC-3’

ITGB1: Forward: 5’-CCTGAGAGTGATGCTACTCCA-3’

Reverse: 5’-CACCCTGGTTGTGCCAAAAAT-3’

PCR amplification was carried out on an Agilent AriaMx Real-Time PCR System (Agilent Technologies, Santa Clara, CA, USA). Gene expression levels were normalized to the housekeeping gene β-actin and analyzed using the 2^^−ΔΔCT^ method.

### Statistic

Data were presented as means ± standard deviation (SD), and n indicates the numbers of animals or experiments in each group. Statistical analyses were performed using GraphPad Prism version 9. Student’s t-test were used for two group s comparation, one-way ANOVA or two-way ANOVA were used to compare more than two groups. P values are described in the legend for each figure, and p-value of less than 0.05 was considered statistically significant.

## Results

### DHT delayed hair regrowth both in female and male mice

In order to clarify the role of DHT on hair growth, we established the DHT-induced alopecia mouse model in both female and male mice. Skin color scoring system was adopted based on a published methodology[31], which was shown in Fig 1B. In female mice, the differences became apparent around 8 days post-depilation, and most pronounced changes of skin color and hair regrowth rate between two groups were observed at day 10 after depilation (Fig 1A), while in male mice, the significant differences showed 12 days after depilation. (Fig 2A). Hair regrowth assessment revealed that DHT effectively suppressed the hair regrowth rate around 35% in female mice (Fig 1C), and inhibitory trend was also observed in male mice, which exhibited approximately 23% decrease (Fig 2B). Furthermore, the skin color scores corroborated the inhibitory effect of DHT on hair regrowth in both sexes (Fig 1D and 2C).

**Figure 1.**
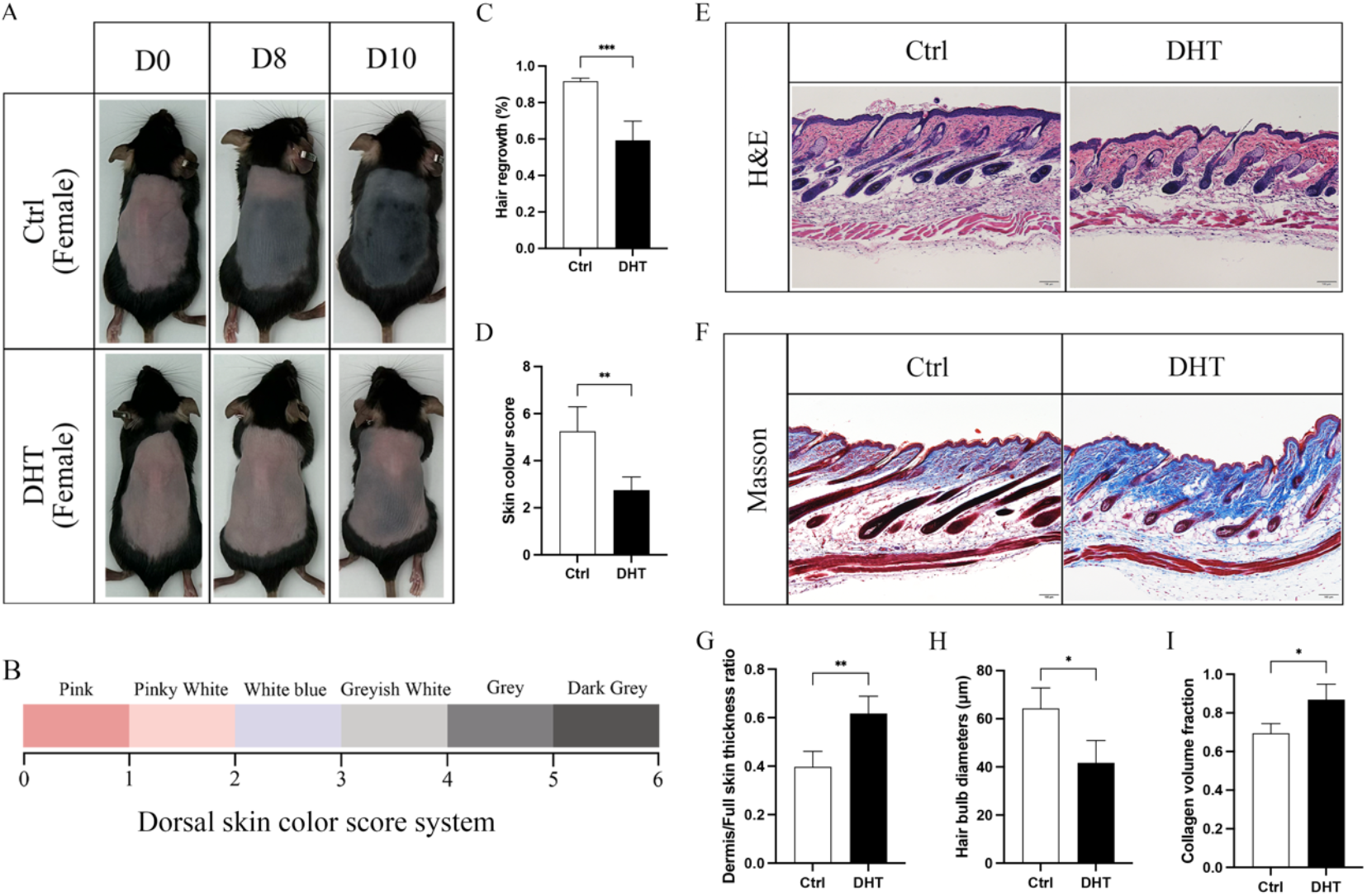
DHT delayed the hair regrowth process in female mice. (A) Dorsal hair regeneration process of female mice treated with Vehicle (Ctrl) or DHT. (B) Dorsal skin color score evaluation system. (C) Hair regrowth rate and (D) skin color score at day 10 were analyzed. (E) Histomorphologic images and (F) Masson trichrome staining images of dorsal skin of mice treated with Vehicle and DHT. Scale bars, 100 μm. DHT altered (G) skin thickness, (H) hair bulb size, and (I) collagen volume fraction. Data were expressed as mean ± S.D. n=4, *P < 0.05, **P < 0.01, ***P < 0.001 compared with ctrl group.

**Figure 2.**
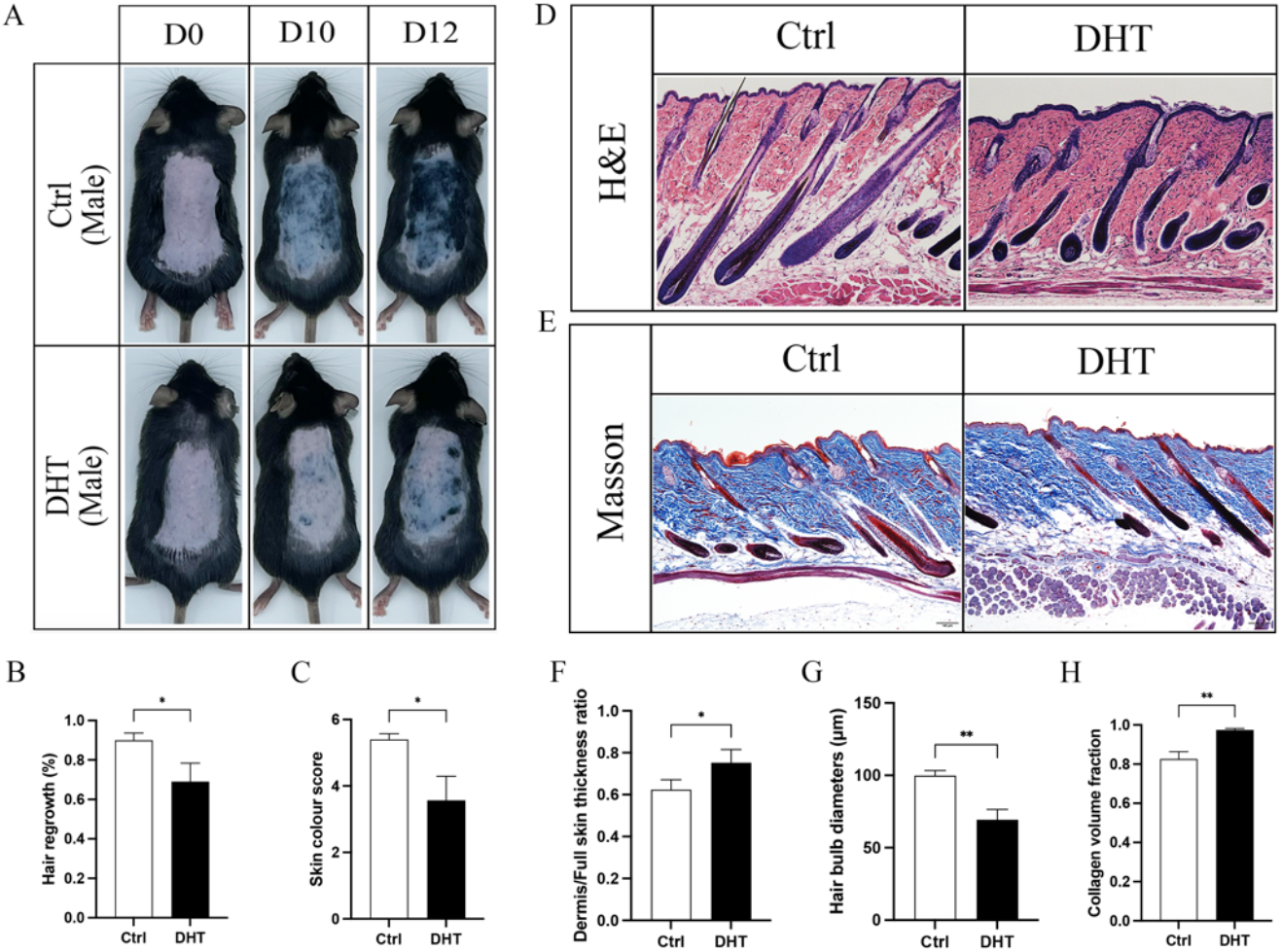
DHT inhibited the hair regrowth process in male mice. (A) Dorsal hair regeneration process of male mice treated with or without DHT. (B) Hair regrowth rate and (C) skin color score at day10 were analyzed. (D) Histomorphologic images and (E) Masson trichrome staining images of dorsal skin of male mice treated with or without DHT. Scale bars, 100 μm. DHT treatment altered (F) skin thickness, (G) hair bulb size, and (H) collagen volume fraction. Data were expressed as mean ± S.D. n=3, *P < 0.05; **P < 0.01 compared with ctrl group.

Histological examination of skin morphology demonstrated that DHT treatment could increase the ratio of dermis to full skin thickness (Fig 1G and 2F) and reduce the hair follicle diameters both in female and male mice (Fig 1H and 2G). Specifically, the dermal thickness ratio of female skin was elevated more than 1.5-fold, and the hair bulb diameters reduced by 35% compared to control group. While in male mice, there were about 1.2-fold increase of dermal thickness ratio, and around 30% reduction of hair bulb diameters compared to control group. Masson’s trichrome staining indicated an increased proportion of collagen volume fraction in DHT-treated mice relative to controls in both sexes (Fig 1I and 2H).

### DHT stimulates AR expression in hair follicle fibroblasts

By using single-cell RNA sequencing technology, we investigated the effects of DHT on various cell types in mouse skin. The dorsal skin samples were collected from both female and male mice of experimental groups, followed by cell isolation and single-cell transcriptome library construction. (Fig 3A). We firstly identified fibroblast cells, which was one of the major cell types (Fig 3B and 3C), with notably high expression of the AR in this population (Fig 3D). GO enrichment analysis of differentially expressed genes (DEGs) in fibroblasts from female mice revealed that DHT administration induced pathways related to skin development, hair follicle morphogenesis, extracellular structure or matrix organization, collagen metabolism and epithelial cell differentiation (Fig 3E). In contrast, in male mice, DHT predominantly enriched pathways including epithelial cell proliferation, extracellular matrix organization, and extracellular matrix organization (Fig 3F).

**Figure 3.**
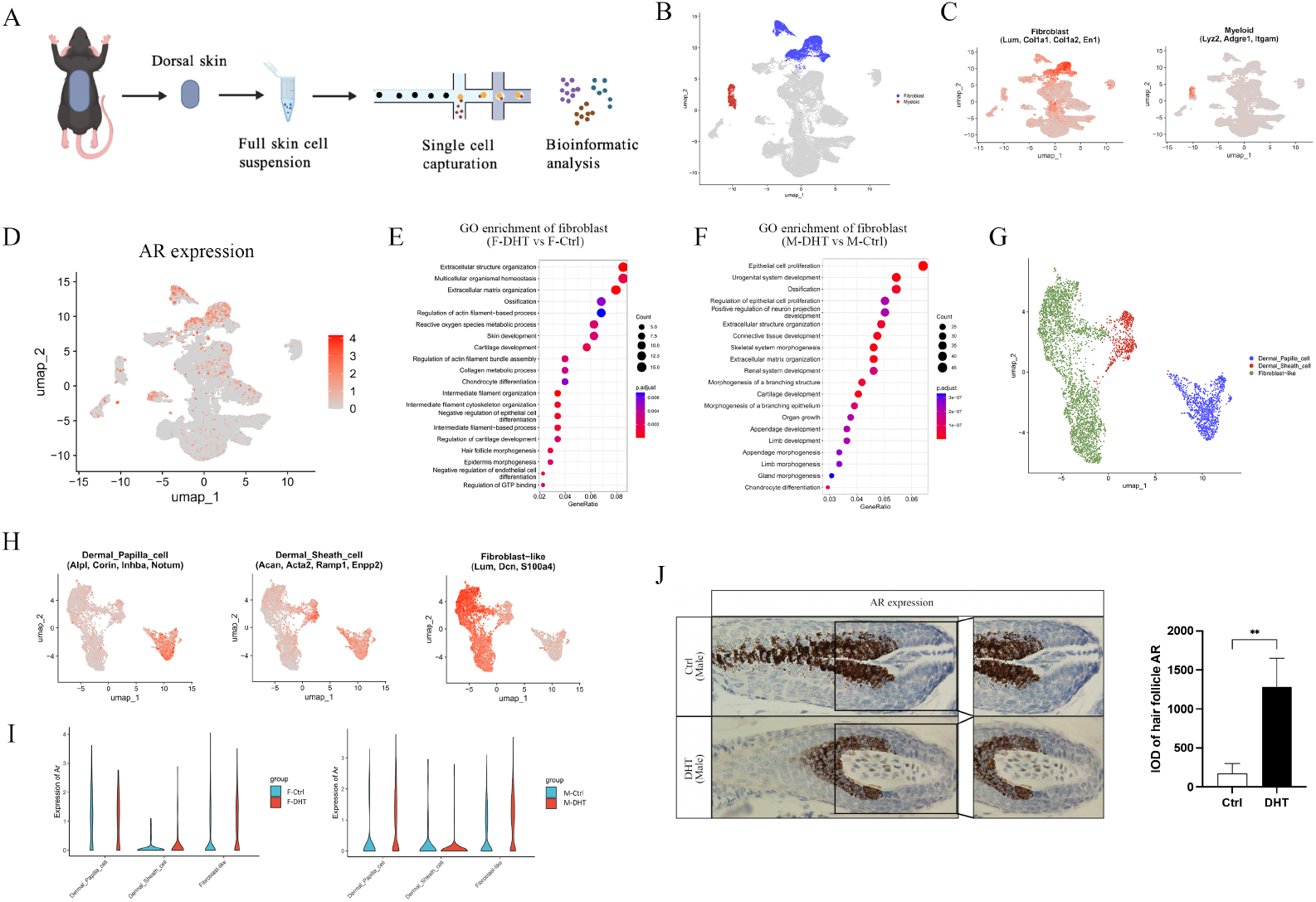
Skin population and AR expression that activated by DHT. We populated skin cells by scRNA sequence. (A) Overview of the experimental workflow, which was created with BioGDP.com[32]. (B) uMAP visualization of skin fibroblast and myeloid cells, colored according to each cell class, annotated by (C) fibroblast and myeloid cell markers. (D) AR expression in skin cells. GO enrichment analysis of DEGs in fibroblast of (E) female mice and (F) male mice. (G) uMAP visualization of fibroblast subclusters, (H) annotated by cell markers. (I) AR expression in fibroblast sub-population. (J) Representative immunohistochemical images showing AR expression by dorsal skin samples of male mice 12 days after depilation treated with or without DHT. Scale bars, 20 μm, and quantification of IOD of AR. Data were expressed as mean ± S.D. n=3, **P < 0.01 compared with ctrl group.

Subclustering of fibroblasts further defined three main subpopulations, including DPC, dermal sheath cells (DSCs), and fibroblast-like cell cluster (Fig 3G and 3H). DHT-mediated upregulation of AR was primarily observed in DPCs and fibroblast-like cells, rather than in DSCs, in both female and male mice skin (Fig 3I). Immunohistochemical analysis of male mouse hair follicles showed abundant AR expression in the bulge region, which corresponds to the location of DPCs. Furthermore, DHT treatment enhanced AR expression by 7-fold increase in this area (Fig 3J), indicating that DHT stimulation activates AR expression within mouse hair follicles.

### Sexually dimorphic apoptosis and immune regulation in DHT-treated skin

GO enrichment analysis revealed that apoptotic signaling pathways were significantly enriched in DPCs, but not in DSCs in DHT-treated female mice (Fig 4A). In DHT-treated male mice, pathways, such as epithelial cell activities, Wnt signaling, mesenchymal activity, and extracellular organization were enriched in both DPCs and DSCs (Fig 4B). Further analysis of apoptosis-related pathways showed that DPCs exhibited enriched apoptotic activities in both DHT-treated female and male mice (Fig 4C and 4D). In contrast, significant enrichment of apoptotic pathways was observed only in male DSCs cells following DHT treatment (Fig 4D).

**Figure 4.**
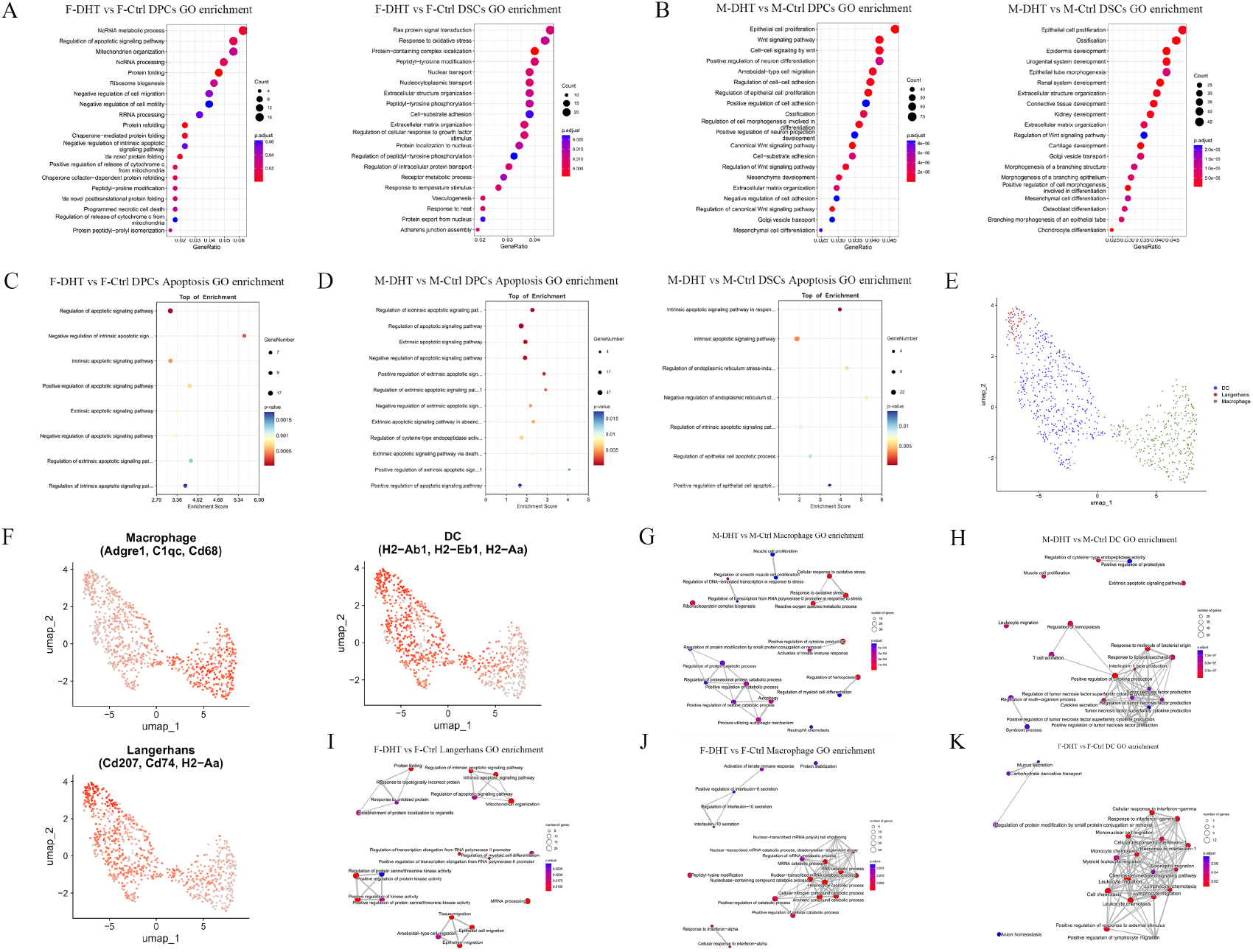
Subpopulation of skin fibroblast and immune cells. GO enrichment analysis of DEGs in DPCs and DSCs in (A) female mice and (B) male mice. Apoptosis-relative pathways analysis of DEGs in DPCs and DSCs in (C) female mice and (D) male mice. (E) uMAP visualization of immune cell subpopulation. (F) Cell annotation of immune cell subclusters. GO enrichment emapplot of DEGs in (G) macrophage and (H) DC in male mice. GO enrichment emapplot of DEGs in (I) LC, (J) macrophage, and (K) DC in female mice.

Given the reported role of immune cells in DHT-induced alopecia [23], we further focused on skin-resident myeloid cells. Myeloid cells were identified (Fig 3B and 3C) and subclustered into three main subpopulations, which were dendritic cells (DCs), Langerhans cells (LC), and macrophages (Fig 4E and 4F). Emapplot analysis of GO enrichment in female macrophages revealed three primary functional modules, including an immune module, a catabolic process module, and an interferon-alpha module, the last was spatially distinct from the main immune-related cluster and was thus considered an independent module (Fig 4J). In male mice, macrophage exhibited greater complexity, with additional modules such as oxidative stress, and ribonucleoprotein complex biogenesis were observed, alongside immune and catabolic process modules (Fig 4G). GO enrichment in female DCs indicated that the immune module was the most responsive to DHT treatment, including leukocyte, lymphocyte, chemokine, and interleukin signaling (Fig 4K). Although an immune module also predominated in males, the specific pathways differed from those in females, with a stronger emphasis on the tumor necrosis factor production. leukocyte migration was regarded as an independent entity due to its separation from the core immune module. Additionally, peptidase activity and extrinsic apoptotic signaling pathway were also observed (Fig 4H). Furthermore, enrichment analysis of LC in females highlighted four key modules, including apoptotic signaling, protein folding, kinase activity, and epithelial cell activities (Fig 4I). No significant enrichment was detected in the corresponding male cell type, suggesting a sex-specific functional activation of these cells.

### Differential interactions between myeloid and fibroblast subpopulations in response to DHT

To investigate potential interactions between the three myeloid subpopulations and the three fibroblast subclusters, we analyzed the number and strength of cell–cell communications inferred from scRNA-seq data. We found the number of interactions in females increased to 1295 after DHT treatment (Fig S1A), whereas males exhibited only 492 interactions despite a post-treatment increase (Fig S1C). An enhancing trend in interaction strength was also observed in both sexes (Fig S1B and 1D). The specific interaction patterns between myeloid and fibroblast subpopulations are summarized in Fig 5A and 5C. In DHT-treated female mice, interactions between DCs and DPCs decreased slightly in both number and strength. In contrast, interactions between DCs and DSCs increased in number but decreased in strength. Interactions between LC and DPCs were enhanced in both quantity and intensity. The interaction of and were enhanced not only in numbers, but also in strength. Conversely, it was found reductions in both the number and strength of interactions between macrophages and DPCs. Although interactions between macrophages and DSCs increased slightly in number, their strength remained largely unchanged (Fig 5A). In male mice, DHT enhanced the number and strength of interactions between DCs and DPCs, while diminished those between DCs and DSCs. Interactions between macrophages and DPCs were strengthened, whereas those between macrophages and DSCs were weakened (Fig 5C).

**Figure 5.**
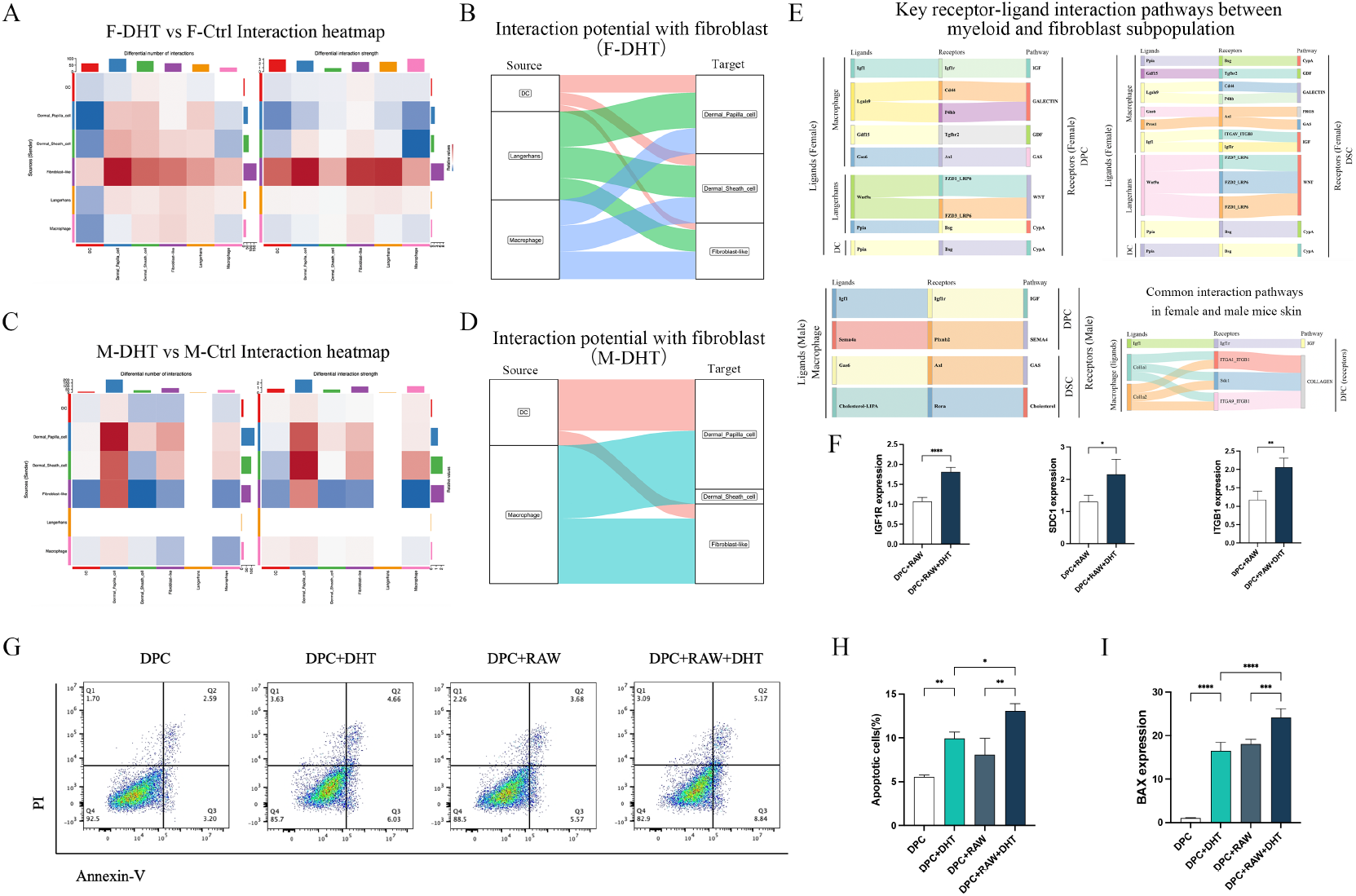
The interactions between fibroblast and myeloid cells. Interaction heatmap of each subclusters in (A) female and (C) male mice. Interaction potential with fibroblast after DHT administration in (B) female and (D) male mice. (E) Key receptor-ligand interaction pathways between myeloid and fibroblast subpopulation in female and male mice. DPCs were co-cultured with RAW 264.7 and stimulated with DHT for 24h, (F) IGF1R, SDC1, and ITGB1 expression in DPCs were examined by qPCR. In indicated experiment, DPCs were collected and stained with Annexin V-FITC and PI, then tested by Flow cytometer. (G) Apoptotic cells proportion when DPCs treated with DHT, with or without RAW264.7. (H) Quantification of apoptotic cells of (G). (I) BAX expression in DPCs treated with DHT, with or without RAW264.7 examined by qPCR. Data are expressed as mean ± S.D. n=4, *P < 0.05, **P < 0.01, ***P < 0.001, ****P < 0.0001.

Based on receptor-ligand pairing possibilities between myeloid and fibroblast subpopulations, Sankey diagram visualization revealed distinct pairing patterns in female and male mice following DHT treatment. In female mice, DPCs and DSCs exhibited a higher number of receptor-ligand pairing possibilities, and predominantly formed with LC and macrophages (Fig 5B). In contrast, in DHT-induced male mice, got more paired possibilities were observed in DPCs and Fibroblast-like cells, largely formed by DCs and macrophages. DSCs had the most pairs formed by macrophage, while no significant pairing was detected involving LC (Fig 5D). In female mice, some of the key pathways between myeloid and DPCs subpopulations included IGF, Galectin, GDF, GAS, Wnt and CypA signaling. Interactions between myeloid cells and DSCs involved similar pathways except for Pros signaling, though often through different receptors within the same pathway. Interestingly, in DHT-stimulated male skin, only a limited set of pathways were engaged, primarily through interactions with macrophages. Key pathways between macrophages and DPCs included IGF and Sema4a, while interactions between macrophages and DSCs were mediated by GAS and Cholesterol pathways. The common pathways between macrophages and DPCs in both sexes were IGF and Collagen signaling (Fig 5E). To functionally validate these findings, we co-cultured human DPCs (hDPCs) with RAW264.7 and examined the expression of receptors associated with the common pathways. It was found that DHT up-regulated the expression of IGF1R, SDC1, and ITGB1 in DPCs at the present of macrophages (Fig 5F).

Given the enrichment of multiple apoptosis pathways in DPCs in both sexes, along with the common interaction pathways between macrophages and DPCs, we further investigated the functional crosstalk by using co-cultured experiment that mentioned above. Flow cytometry results showed that DHT alone increased the percentage of apoptotic hDPCs, and this effect was further enhanced when hDPCs were co-cultured with RAW264.7 (Fig 5G and 5H). This pro-apoptosis effect was corroborated by qPCR, which exhibited a marked upregulation of the BAX expression following DHT stimulation, and was further amplified at the present of RAW264.7 (Fig 5I), suggesting a synergistic role of macrophages in promoting DHT-induced apoptosis in DPCs.

### Y001 and Y002 intervention could promote hair regrowth in mice

Y001 and Y002 are two types of novel polypeptides that exhibit hair regrowth-promoting properties. Based on our previous finding that DHT induces apoptosis in DPCs, we first evaluated the effects of Y001 and Y002 on DHT-mediated apoptosis of DPCs. Flow cytometric analysis demonstrated that both polypeptides significantly attenuated DHT-induced early apoptosis in DPCs (Fig 6A and 6B). We next applied Y001 and Y002 topically in DHT-induced alopecia models in both female and male mice. Both compounds markedly alleviated DHT-induced suppression of hair regrowth. Significant improvements were clearly observed at day 13 post-depilation (Fig 6C and 6E), with quantitative analyses indicating that Y001 and Y002 enhanced the hair regrowth rate by more than 1.3 folds in female and 1.2 folds in male mice when compared to DHT group (Fig 6D and 6F). Histological examination of dorsal skin samples revealed that DHT significantly reduced hair bulb diameters in both sexes, whereas treatment with Y001 or Y002 increased bulb diameters by approximately 1.6 folds in female, and 1.3 folds in male mice (Fig 6G-6J).

**Figure 6.**
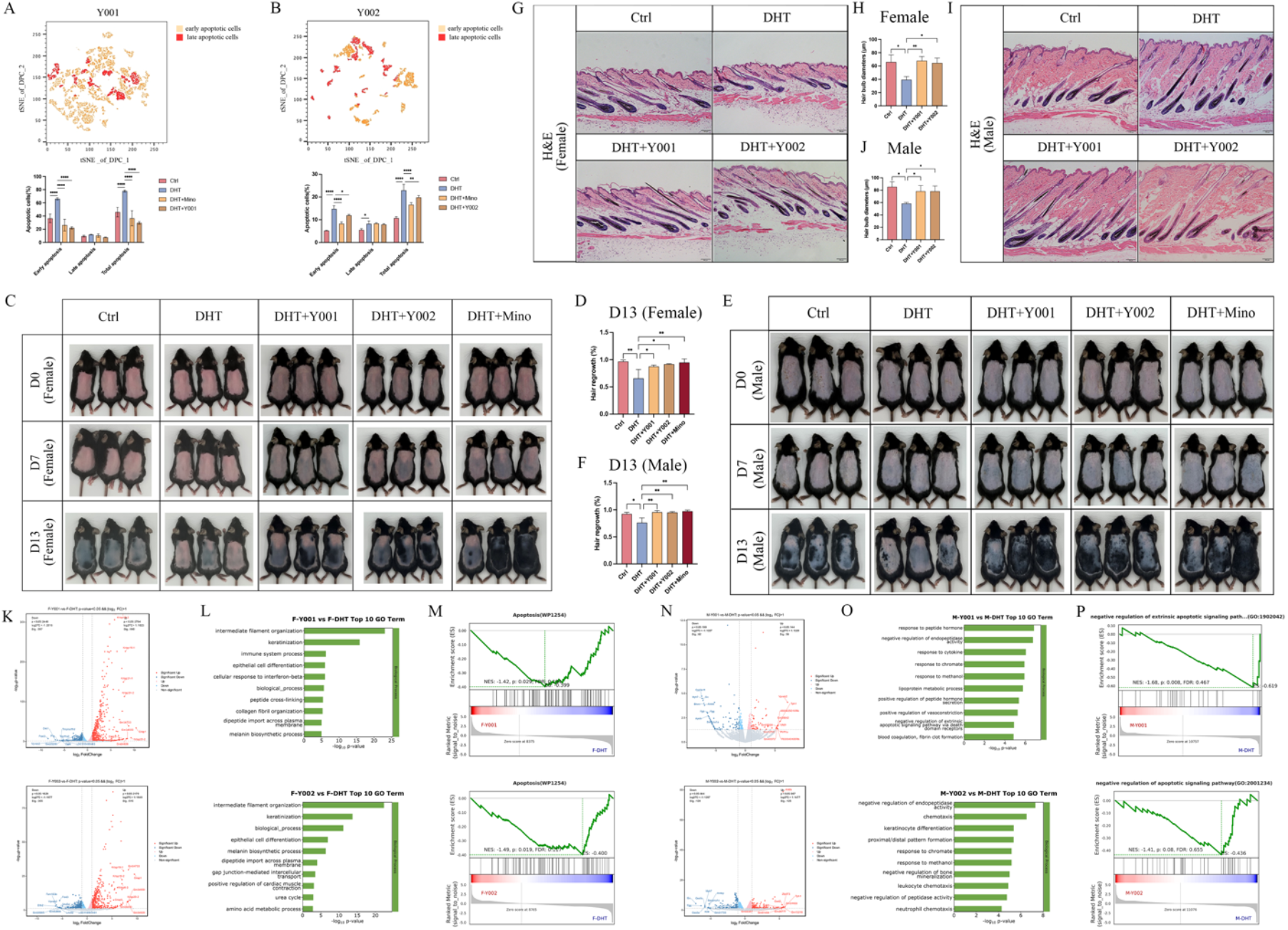
Y001 and Y002 could promote the hair regrowth in DHT-induced alopecia mice models. Y001 and Y002 were applied to DPCs treated with DHT for 48h. DPCs were collected and stained with Annexin V-FITC and PI, then tested by Flow cytometer. Apoptotic cells proportion after (A) Y001 and (B) Y002 application, and quantification of apoptotic cells. Representative images of Y001 and Y002 that applied in (C) female or (E) male mice dorsal skin, respectively, minoxidil was the positive control. Quantification of (D) female or (F) male hair regrowth rate at day 13 after depilation. Histomorphologic images of dorsal skin of (G) female or (I) male mice treated with or without DHT, Y001 and Y002. Scale bars, 100 μm. Quantification of hair bulb size of each group in (H) female and (J) male mice, respectively. Volcano plot of DEGs between Y001 or Y002 and DHT group in (K) female and (N) male mice. GO enrichment analysis of DEGs between Y001 or Y002 and DHT group in (L) female and (O) male mice. GESA enrichment of apoptosis pathway in Y001 and Y002 treated (M) female and (P) male skin. Data are expressed as mean ± S.D. n=3, *P < 0.05; **P < 0.01, ****P < 0.0001.

To explore the underlying mechanisms, we conducted bulk RNA sequencing on dorsal skin samples from each experimental group and performed functional enrichment analysis to identify transcriptomic changes. In female mice, compared to DHT group, treatment with Y001 significantly upregulated 665 genes and downregulated 587 genes, while treatment with Y002 increased 515 genes and reduced 355 gene (Fig 6K). GO enrichment analysis showed that intermediate filament organization, keratinization, epithelial cell differentiation, and melanin biosynthetic process were enriched in both treatment groups, besides, immune response was also enriched after Y001 administration (Fig 6L). Both Y001 and Y002 downregulated DHT-induced apoptotic pathway (Fig 6M) and reversed several DHT-upregulated pathways, including immune system process and collagen fibril organization. Furthermore, several pathways downregulated by DHT, including keratinization, epithelial cell differentiation, and melanin biosynthetic process were restored by both treatment (Fig S2A-B). In male mice, Y001 administration significantly upregulated only 58 genes and downregulated 82 genes, whereas Y002 treatment resulted in a comparable number of upregulated and downregulated genes, which were 125 and 128, respectively (Fig 6N). GO analysis indicated that Y001 application enriched pathways including peptide hormone, response to cytokine, and extrinsic apoptotic signaling, while Y002 treatment enriched chemotaxis, keratinocyte differentiation, and peptidase activity (Fig 6O). Notably, negative regulation of extrinsic apoptotic pathway and negative regulation of apoptotic pathway were downregulated by Y001 and Y002, respectively (Fig 6P). However, none of the top 10 DHT-downregulated pathways were reversed in males. Instead, Y001 downregulated several DHT-induced pathways, such as response to peptide hormone, lipoprotein metabolic process, cholesterol efflux, and negative regulation of endopeptidase activity (Fig S2C). Y002 also counteracted DHT effects by modulating pathways such as negative regulation of endopeptidase activity, cardiac muscle contraction, and response to peptide hormone (Fig S2D). These findings suggest that Y001 and Y002 mitigate DHT-induced inhibition of hair regrowth not only through a suppression of apoptotic signaling, but also distinct sex-specific mechanisms.

## Discussions

The androgenetic alopecia is primarily driven by androgens, such as DHT, however, its pathogenesis also involves complex interactions between the skin immune microenvironment and hair follicle cells. Our study elucidates how myeloid cells interact with fibroblasts in both female and male mice, providing insight into the sexual dimorphism underlying DHT-induced hair follicle regression. Furthermore, we demonstrate that two polypeptides, Y001 and Y002, promote hair regrowth in a DHT-mediated mouse model of alopecia, suggesting their potential as therapeutic strategies for androgenetic alopecia.

We first confirmed that systematic administration of DHT delayed hair regrowth process both in female and male mice, which is consistent with previously reported effects in male mice[31]. Increasing evidence suggests an important role of inflammation in the pathogenesis of AGA[33, 34]. Notably, certain anti-inflammatory treatments have yielded therapeutic outcomes. For example, cell-free fat extract, derived from adipose tissue and rich in anti-inflammatory factors, has been shown to promote the hair growth in AGA patients, likely mediated through its anti-inflammatory components[35]. Although inflammatory markers are not typically detected in plucked hair follicles, their elevated expression has been observed in the infundibulum and scalp skin of individuals with AGA[36]. By using scRNA-seq technology, we characterized the cellular landscape of DHT-exposed skin and identified the subpopulations of skin fibroblast and myeloid cells. LC are unique tissue-resident macrophages with DC functionality, and mainly reside in the epidermis[37]. Interestingly, we found although LC were present in the skin of both sexes, no significant functional enrichment of DEGs was detected in male mice. Enrichment analysis further highlighted immune response as a prominent characteristic of macrophage and DCs in both female and male mice. This sexual dimorphism was particularly evident in the activation patterns of apoptotic pathways, as multiple apoptotic signaling pathways enriched in DPCs in both sexes, which is in line with previous reports[26], while apoptotic pathways enriched in DSCs observed exclusively in male mice.

Furthermore, we employ scRNA-seq to reveal, for the first time, the sexual dimorphism in immune microenvironment-hair follicle cell interactions in androgenetic alopecia, offering new perspectives on the mechanisms underlying androgen-driven hair loss. we found that DHT enhanced interactions of myeloid and fibroblast subpopulations in both female and male mice, but females exhibit more robust interactome and pathway richness. For example, a more complex signaling network between macrophages and DPCs or DSCs were observed in female mice. Additional signaling molecules, such as FGF10, which has been reported in previous study, may also contribute to the interplay of macrophage and DPCs[38]. Numerous interaction pairs involving LC were observed in female mice than in male mice, suggesting a sex-specific role for these cells in DHT-mediated skin. However, only two pathways were commonly shared between macrophages and DPCs in both sexes, including IGF and collagen signaling pathways, mediated by IGF1R, ITGB1 and SDC1. IGF1R involved in the hair follicle regeneration[39, 40] and the transition of anagen to catagen phase[41], potentially by promoting angiogenesis to regulate hair growth[42]. We found DHT treatment upregulates the expression of IGF1R in DPCs, this is consistent with the studies of reported prostate cancer that androgen-stimulated AR signaling could enhance IGF1R transcription [43, 44], suggesting a similar interplay may occur during hair loss progression. Although AR overactivation is known to trigger cascade signaling and finally may induce the hair follicle apoptosis[45], whether interactions between IGF1 and androgen axes contributes to this process requires further investigation. The role of SDC1 in hair follicles remains poorly understood. It is dynamically expressed during the hair cycle, predominantly in the anagen outer root sheath and dermal papilla[46]. We found an upregulation expression of SDC1 in DPCs, which may imply a potential role in modulating microenvironmental signaling, though the specific molecular mechanisms remain to be elucidated. Enrichment analysis indicated extracellular matrix (ECM) activities, potentially regulated through ITGB1, which was reported for regulating signals when ECM bind to the extracellular region[47]. This suggests that DHT may influence follicle regression by altering cell–matrix interactions. In vitro experiments supported a synergistic effect of macrophages and DHT on DPCs.

Finally, we evaluated two polypeptides, Y001 and Y002 in DHT-induced alopecia mouse model. Both peptides significantly promoted hair regrowth in both female and male mice. The underlying mechanism may involve anti-apoptosis effects on DPCs under DHT stimulation. Furthermore, administration of Y001 and Y002 reversed several DHT-induced signaling pathways. In female mice, DHT-induced upregulation of immune response was reversed by Y001 and Y002 application, suggesting a modulation of immune cell activities. In male mice, both peptides appilcation enriched immune-related pathways, however, the reversed regulatory effects were more prominent in hormone-related pathways. The efficacy observed across sexes highlights the clinical translational potential of Y001 and Y002 as a universal therapeutic strategy for androgenetic alopecia regardless of gender.

In conclusion, our study highlights an essential contribution of the immune microenvironment to androgen-mediated alopecia, and reveals its distinct gender-dependent effects. Furthermore, we evaluate polypeptides Y001 and Y002 as highly promising candidates for treatment due to their ability to promote hair regrowth and mitigate DHT-induced apoptosis in DPCs. Future work will focus on elucidating the precise molecular targets of these peptides and validating their efficacy and safety in advanced preclinical models, paving the way for novel treatments for androgenetic alopecia.

## Abbreviation

AGA: Androgenetic alopeica
FPHL: female pattern hair loss
DHT: Dihydrotestosterone
AR: Androgen receptor
scRNA-seq: single-cell RNA sequencing
DPCs: dermal papilla cells
DSCs: dermal sheath cells

## Author Contributions

Conceptualization, J.V. Z.; methodology, FF D., L G., YL Y., JM W., ZC S., J C., MX L. and DF Z.; validation, FF D, L G. and YL Yang.; formal analysis, FF D, L G. and YL Y.; investigation, resources, data curation, FF D. and TX X., writing—original draft preparation, FF D.; writing—review and editing, FF D., L G., YL Y., JM W., ZC S., J C., MX L., TX X., DF Z. and J.V.Z., visualization, FF D., L G., YL Y. and DF Z., supervision, J.V. Z., DF Z., TX X., and ZL D.; project administration, FF D.; funding acquisition, J.V. Z. and ZL D. All authors agreed to the published version of the manuscript.

## Acknowledgments

This study was supported by the National Key R&D Program of China (2024YFA1803001), Shenzhen Medical Research Fund (B2404004), National Natural Science Foundation of China (32571489, 82301907, 32502180), Guangdong Basic and Applied Basic Research Foundation (2024A1515010059, 2024A1515030279), Shenzhen Science and Technology Program (JCYJ20220818101218040, JCYJ20220818102811025, JCYJ20220818103608017, JCYJ20220818103607015, JCYJ20230807140805011, JCYJ20220531095811025), Shenzhen Key Laboratory of Metabolic Health (Grant No. ZDSYS20210427152400001), National Natural Science Funds for Excellent Young Scientists (No. 82422063), National Key Research and Development Program of China (No. 2023YFC2509003).

