## Supplementary figures and images for "Crosstalk between Immune Microenvironment and Hair Follicle Cells Underlies Sexual Dimorphism in Androgenetic Alopecia"

### Fig S1-1.tif

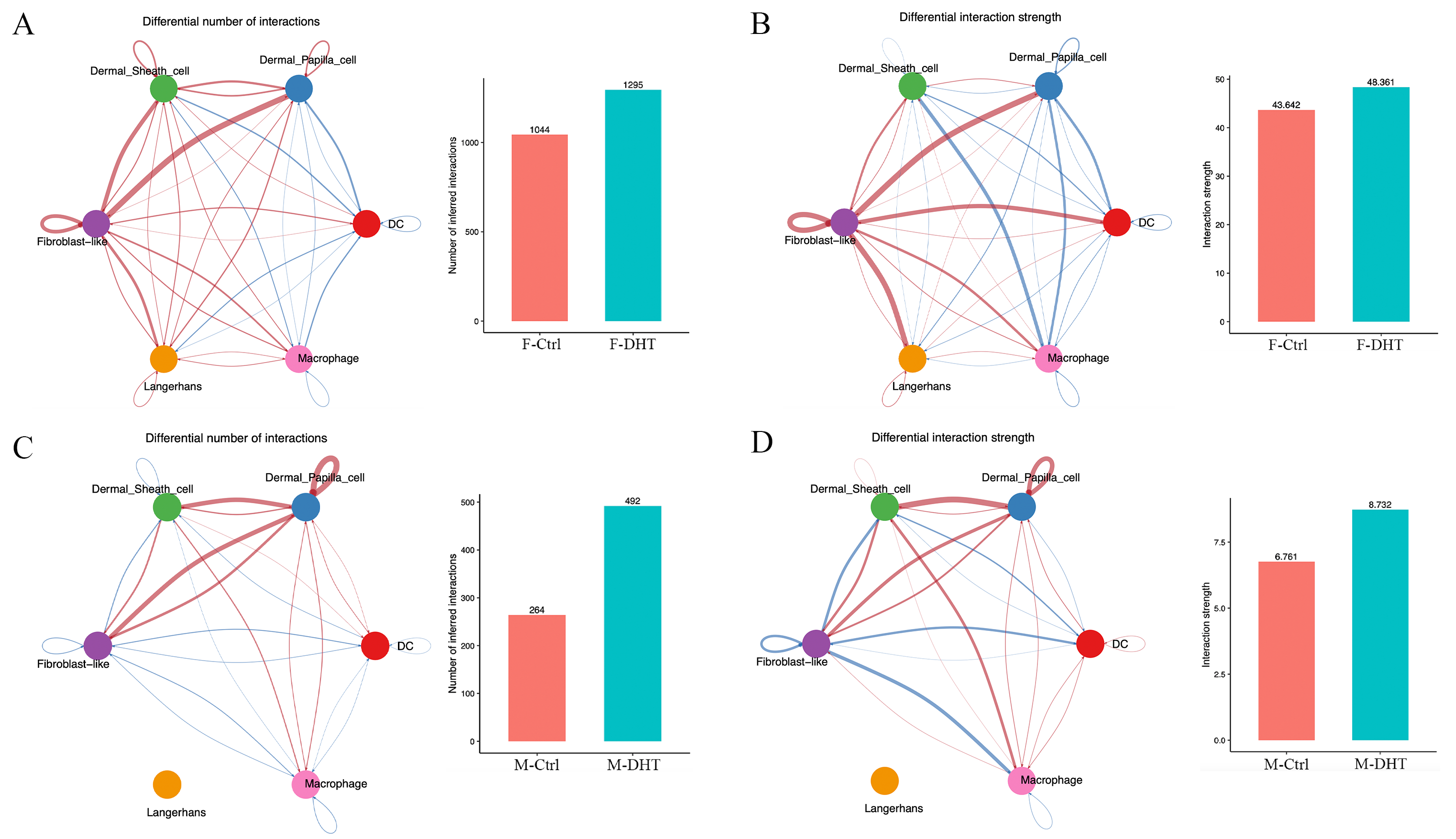

### Fig S2-1.tif

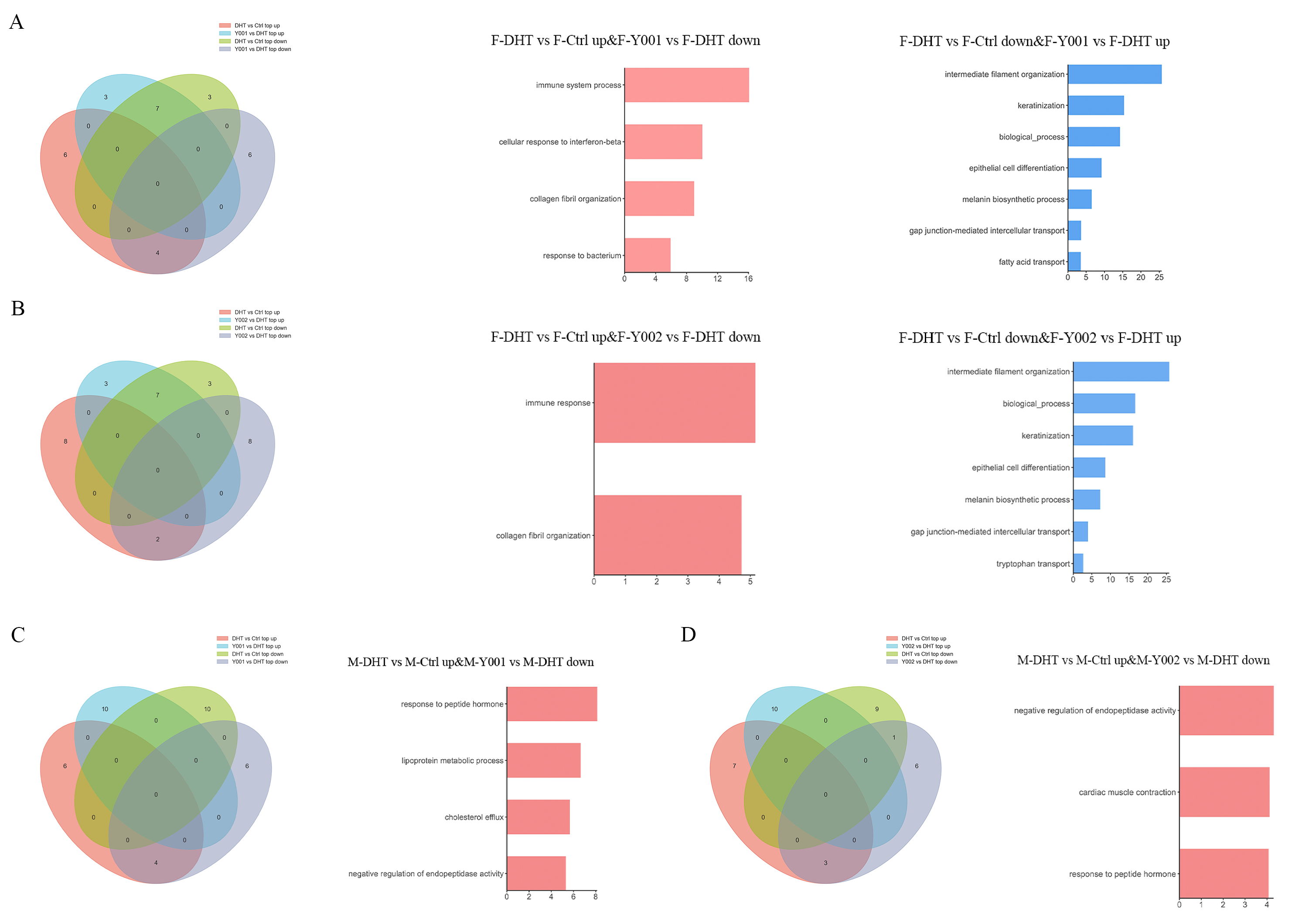
